# Realized generation times: contraction and impact of infectious period, reproduction number and population size

**DOI:** 10.1101/568485

**Authors:** Andrea Torneri, Amin Azmon, Christel Faes, Eben Kenah, Gianpaolo Scalia Tomba, Jacco Wallinga, Niel Hens

## Abstract

One of the key characteristics of the transmission dynamics of infectious diseases is the *generation time* which refers to the time interval between the infection of a secondary case and the infection of its infector. The generation time distribution together with the reproduction number determines the rate at which an infection spreads in a population. When defining the generation time distribution at a calendar time *t* two definitions are plausible according whether we regard *t* as the infection time of the infector or the infection time of the infectee. The resulting measurements are respectively called *forward generation time* and *backward generation time*. It has been observed that the mean forward generation time contracts around the peak of an epidemic. This contraction effect has previously been attributed to either competition among potential infectors or depletion of susceptibles in the population. The first explanation requires many infectives for contraction to occur whereas the latter explanation suggests that contraction occurs even when there are few infectives. With a simulation study we show that both competition and depletion cause the mean forward generation time to contract. Our results also reveal that the distribution of the infectious period and the reproduction number have a strong effect on the size and timing of the contraction, as well as on the mean value of the generation time in both forward and backward scheme.

**Author summary:** Infectious diseases remain one of the greatest threats to human health and commerce, and the analysis of epidemic data is one of the most important applications of statistics in public health. Thus, having reliable estimates of fundamental infectious diseases parameters is critical for public health decision-makers in order to take appropriate actions for the global prevention and management of outbreaks and other health emergencies. A key example is given by the prediction models of the reproduction numbers: these rely on the generation time distribution that is usually estimated from contact tracing data collected at a precise calendar time. The forward scheme is used in such a prediction model and the knowledge of its evolution over time is crucial to correctly estimate the parameters of interest. It is therefore important to characterize the causes that lead to the contraction of the mean forward generation time during the course of an outbreak.

In this paper, we firstly identify the impact of the epidemiological quantities as reproduction number, infectious period and population size on the mean forward and backward generation time. Moreover, we analyze the phenomena of competition among infectives and depletion of susceptible individuals highlighting their effects on the contraction of the mean forward generation time. The upshot of this investigation is that the variance of the infectious period distribution and the reproduction number have a strong impact on the generation times affecting both the mean value and the evolution over time. Furthermore, competition and depletion can both cause contraction even for small values of the reproduction number suggesting that, in epidemic models where the generation time is considered time-inhomogeneous, estimators accounting for both depletion and competing risks are to be preferred in the inference of the generation interval distributions.

## Introduction

In infectious disease epidemiology, mathematical models are increasingly being used to study the transmission dynamics of infectious agents in a population and thereby providing fundamental tools for developing control policies. An optimal control strategy is based on an appropriate prediction model that in turn requires reliable estimates of the key epidemic parameters.

Most research has focused on the ‘basic reproduction number’, *ℛ*_0_, which is defined as the expected number of secondary cases resulting from introducing a typical infected person into an entirely susceptible population [2]. The inference of its value in the ascending phase of an epidemic is based either explicitly or implicitly on assumptions about the generation interval distribution [3].

The generation interval, or generation time, is defined to be the time interval between the infection time of an infectee and the infection time of its infector [4]. Generation times are lengths of time intervals and thus there is not a unequivocal procedure to define their dependence on a precise calendar time *t*. To account for the evolution over time a choice has to be made weather considering generations from the infectee or infector point of view. In the former case the time coordinate refers to the time that has evolved since the infector of an infected person was infected. This is called ‘backward’, or ‘period’, generation interval. In the latter case, known as ‘forward’, or ‘cohort’, generation interval the average time required to infect another individual is recorded [5, 6]. Considered in the forward scheme, the generation interval distributions is commonly used to estimate infectious disease parameters such as the basic reproduction number [6–9].

More ambiguity arises in the estimation of the mean generation time because actual data often concern the onset of clinical symptoms rather than the time of infection. These observations relate to the ‘serial interval’, the time interval between symptom onset of a secondary case and symptom onset of its infector [10]. Many authors have used the serial interval as a synonym for the generation time. However, unlike the generation time, the serial interval can have a negative duration when the clinical symptoms for an infectee appear prior to that of its infector [11]. Note that the serial interval is only defined for symptomatic individuals; an issue that we will not discuss here.

Statistical development led to approaches for the estimation of the generation time distribution [7, 12, 13] or jointly of the basic reproduction number and the generation time distribution [14–16]. The usefulness of the aforementioned approaches has been demonstrated in the analysis of epidemic data during e.g. SARS outbreaks and the pandemic influenza A(H1N1)V2009 outbreak [17–19]. Most of these estimation methods assume the generation or serial time distribution to remain constant during the epidemic. However, several authors described a non-constant evolution over time for both backward and forward generation interval [5, 6, 8, 20]. In the former case as the epidemic evolves the generation time increases while in the latter case the generation time contracts reaching a minimum approximately at the peak of the outbreak [6, 8]. We will refer to this phenomenon in the forward scheme as ‘contraction’ to stress the particular shape that the mean generation time assumes over time. The non-constant evolution of the generation interval has stimulated a search for different approaches to estimate the reproduction number that avoid assuming a constant generation intervals distribution through time [8, 9, 13]. Kenah et al. (2008) proposed an hazard-based estimator and the so-called contact interval, the time from onset of infectiousness to an infectious contact, accounting not only for depletion but also for competing risks.

The contraction of the mean forward generation time seems counter-intuitive since one would expect generations to happen faster in the initial phase of the epidemic, when the population is mostly susceptible. The principal aim of this paper is to clarify the epidemiological mechanisms that cause contraction. Researchers typically assign to the phenomenon of contraction only one among two explanations: competition among infectors [8] and depletion of susceptible individuals [5, 20]. Both explanations are reasonable, but a study that clearly shows which of these hypotheses are responsible for the contraction of the mean forward generation time is not present in literature. The first explanation requires multiple infectors competing to infect the same susceptible and affects the specific generation time while the latter accounts for the variation in the probability of encountering a susceptible individual during the outbreak inducing infectors to more likely infect other individuals in a short time frame since the probability of contacting a susceptible later on is lower. More recently, Liu et al. (2018) reported the evolution over time of the mean forward generation interval in a structured population. More precisely, different infectious contact processes have been defined in different locations showing that levels of contraction strongly depend on the underlying contact process. Therefore, in addition to investigate the competition and depletion hypotheses, we address the impact on the mean backward and forward generation intervals over time of settings with different reproduction numbers, population sizes and infectious period distributions.

The present investigation is a simulation study where the simulations are based on two different algorithmic implementations of a stochastic SIR compartmental model as introduced by Kenah et al. (2008) and Scalia Tomba et al. (2009) which will be referred to as the stepwise and the parallel algorithm respectively. In what follows, we revisit the taxonomy of various time intervals used in epidemic modeling, we simulate the impact of the considered epidemic quantities on the mean generation times and we show the impact of competition and depletion on the mean forward generation interval. In addition to a baseline scenario we present three artificial scenarios accounting for different levels of competition and depletion level since in the SIR model these phenomena cannot be disentangle. Finally, we conclude by explaining under which conditions the two hypotheses give rise to the generation interval contraction.

## Materials and methods

We follow the notation used by Kenah et al. (2008) to describe the dynamics of the well-known ‘Susceptible-Infected-Removed’ (SIR) compartmental epidemic model in a closed population of size *N*:

1. after an infectious contact, person *i* acquires infection at time *t*_*i*_.
2. during the infectious period of length *r*_*i*_, person *i* is capable of infecting other individuals
3. at time *t*_*i*_ + *r*_*i*_, person *i* is immune and cannot be longer infected by other person.

We distinguish between three time intervals that determine the between-host transmission of infection. The first one is the *contact interval τ*_*i,j*_, defined to be the time interval between the onset of infectiousness in person *i* and the first infectious contact from *i* to *j*, where we define an infectious contact as a contact sufficient to transmit the disease.

After becoming infectious at time *t*_*i*_ an infected person *i* makes contact with person *j* at time *t_i,j_* = *t*_*i*_ + *τ_i,j_*. When the contact interval *τ_i,j_* occurs within the infectious period *r*_*i*_, i.e. *τ*_*i,j*_ < *r*_*i*_, the infection can be transmitted and the contact interval is called *infectious contact interval* 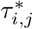. This contact will lead to an infection if *j* is susceptible at time *t*_*i,j*_. In this framework the *generation time ω*_*i,j*_ can be defined in the following way: if an infectious contact from person *i* to person *j* leads to an infection transmission, then 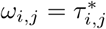 is called the generation time. In the present paper we consider a Poisson contact process.

### Simulation setup

Our simulation models are based on two algorithms previously introduced in literature. The *stepwise* algorithm was proposed by Kenah et al. (2008); it is based on the notion contact interval [22] and it was implemented to illustrate the competition among potential infectors and its effect on the mean generation times. After the onset of infectiousness in an indiviudal, infectious contact intervals are drawn for all the other susceptible persons. The *parallel* algorithm was outlined by Scalia-Tomba et al. (2009); it is based on the infectious contact process aiming to illustrates the effect of the depletion of susceptible individuals. Based on a specific infectious contact process, a single infectious contact interval is proposed by all the individuals that are infectious in the current time step. A complete description of both algorithms is given in S1 Appendix. Since results are similar for both algorithms we report results based on the parallel algorithm and defer results based on the stepwise algorithm to S1 Appendix (S1 Fig and S1 Table).

### Baseline scenario

We start the investigation of the causes that affect the generation time setting a baseline scenario that representing the dynamic of a stochastic SIR model. In the baseline scenario, we look at the impact of the infectious period, the reproduction number and the population size on the mean backward and forward generation interval using the two aforementioned algorithms. In the forward scheme, the mean generation interval is calculated within each infector’s set of generation times and then used as a single data point per infector to avoid the size biased sampling effect [5] whereas in the backward observation scheme, we attribute the unique generation time to each single infectee.

We simulate different epidemics by varying:

- *ℛ*_0_=1.5, 2, 3, 4, 5
- the infectious period distribution with mean 1: constant, Exp(1) = Γ(1, 1), Γ(0.5, 2) and Γ(2, 0.5), resulting in variances equal to 1, 2 and 0.5, respectively.
- the population size: N=100, 500 and 1000

We work with a completely susceptible and closed population and report, based on 1000 simulations, the mean duration of an epidemic (*T*_*max*_), the mean final size (*F*_*s*_), and the averages of mean forward 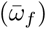 and mean backward generation times 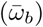. We study non-extinct outbreaks or outbreaks that persist, i.e. outbreaks in which a substantial proportion of the population is infected, i.e. with final size larger than 10% of the total population.

The evolution over time of the generation intervals is reported performing a loess regression on the first *n* = 35 non-extinct simulations. We used this approach to account for the stochasticity of the epidemic process and because we want to attach a confidence interval that quantifies the variability over simulations. The number of simulations is set to *n* = 35 because of computational limitations: the prediction of the loess regression requires a lot of computational memory, especially for high reproduction numbers and high population sizes where a considerable number of generations are registered. Rerunning results on random sets of simulations of size *n* yielded similar results and therefore the choice of *n* = 35 is not considered a limitation. The loess function requires specifying a smoothness parameter called *span*. Different span values did impact the plot in the final part of the epidemic but did not qualitatively affect the results (S3 and S4 Figs). The same approach will be used also for the other scenarios.

### Contraction of the mean forward generation time

We introduce three summary measures to account for competition and depletion. In case the interest is in competition, we compute the relative number of generations affected by competition *p*_*c*_, i.e. the number of generations for which more than an individual propose an infectious contact to a specific susceptible over the total number of generations; we report also the mean number of competitors when there is competition, *µ*_*c*_. In case interest is in depletion, we compute the maximum depletion *ϕ* that is defined to be:

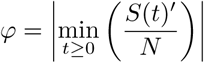

We refer to S1 Appendix for the procedure to calculate the maximum depletion.

We first show that competition and depletion are present in the baseline scenario and we compare these summaries for populations of size *N* = 4, 10, 20, 50, 100, 200 and for *ℛ*_0_ = 1.5, 3, 5.

After that, to investigate the phenomenon of contraction, next to the baseline scenario, we study two scenarios that increase the effect of, respectively, depletion and competition on the generation time distribution. In the former scenario susceptible persons are vaccinated at a specific moment in time during the epidemic, referred to as the *vaccination scenario*, and in the latter scenario infectious individuals are forced to compete for the same susceptible, referred to as the *competition scenario*. We also study a scenario in which the competition among infectors is not present: individual are forced to proposed a contact only to susceptible persons who no one already proposed an infectious contact to. We refer to this scenario as the *pure depletion* scenario. In all of these scenarios the infectious period is set here to be constant to avoid that the stochasticity of the infectious period distribution affects the results.

### Vaccination of susceptible persons

We study simulations in which 30%, 60% and 90% of the susceptible population is vaccinated during the epidemic. In a population of size *N* = 1000 and *ℛ*_0_ = 1.5 we do so by vaccinating the remaining susceptible persons at a specific time called vaccination time and indicated with *t*_*v*_. In this way we change the depletion effect: both augmenting the intensity and changing the time at which depletion occurs. We consider simulations with different vaccination times representing the initial phase (*t*_*v*_ = 2), the main phase (*t*_*v*_ = 3, 5, 7) and the last phase (*t*_*v*_ = 9) of the epidemic and we compute the value of the epidemic characteristics. We do not report the depletion entity because of the instant drop in susceptible population. Lastly, we plot the evolution over time of the forward generation time for comparing this depletion scenario with the baseline scenario.

### Pure depletion scenario

In this scenario infectives propose infectious contacts only to individuals who no other infectors proposed an infectious contact to. In this way, there is no competition and only depletion would be responsible for contraction. We consider populations of size *N* = 100, 500 and 1000 and reproduction numbers *ℛ*_0_ = 1.5, 2, 3, 4, 5 and we compare this scenario with the baseline scenario in which competition is present. For every non-extinct simulation we compute the loess regression and we keep track of the maximum value, the minimum value and the range defined as the difference between maximum and minimum. We do not report here the relative number of generations affected by competition because of requiring huge computational memory to monitor all these generations. However, for a small sample size, values are in line with the simulations for *N* = 100, 200.

### Competition among infectious persons

In the competition scenario, we force individuals to compete for the same susceptible persons in a fixed time frame. To do that, every time an individual proposes an infectious contact we force that individual to propose it to a susceptible person who already someone else proposed an infectious contact to. We consider a scenario with population size *N* = 1000 and *ℛ*_0_ = 1.5 and, without loss of generality, we modify the code increasing competition when the outbreak time is in the interval (3, 7). We select this time interval to allow for starting from a reasonably sized infectious population.

## Results

### Impact of infectious period, reproduction number and population size on the realized generation intervals

Table 1 summarizes the results of the baseline scenario showing remarkable differences when different infectious period distributions are considered: an higher variance of the infectious period distribution enlarges the disparity between the value of the mean forward and backward generation times, specially in the case of low value of the reproduction number. Moreover, the outbreaks last longer and the backward and forward generation times are registered to be larger. This discrepancy between mean backward and forward generation time decreases for scenarios with lower variance resulting almost the same when a constant infectious period is considered. The average mean generation time decreases as the basic reproduction number increases, in line with previous studies [5, 6, 8].

**Table 1.**
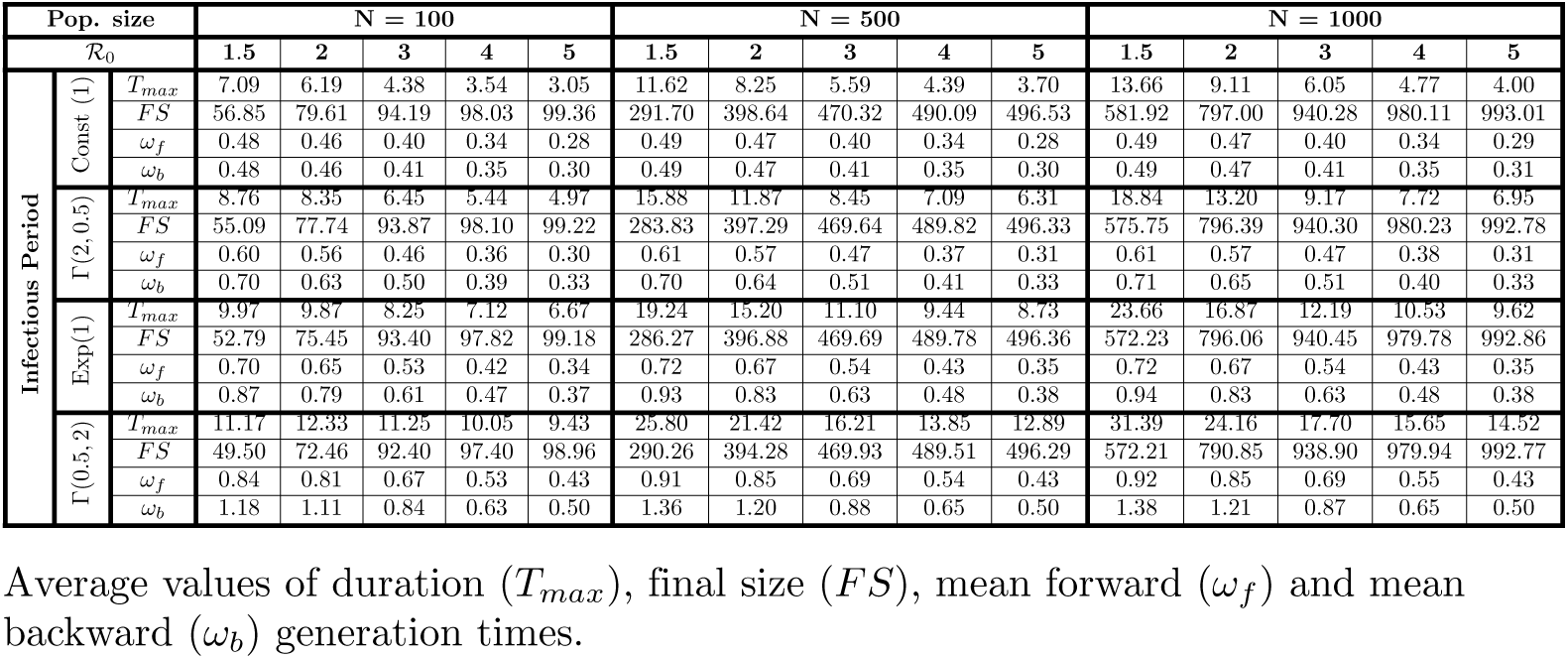
Epidemic characteristics - baseline scenarios

In Figs 1 and 2 we show the evolution over time of the mean forward and backward generation times in a population of size *N* = 1000, respectively, for *ℛ*_0_ = 1.5, 2, 3, 4, 5 and for the infectious period distributions specified before. The mean forward generation interval contracts as the reproduction number increases but still slightly even for low values of the reproduction number. In the backward observation, the generation time shows an increasing trend that is steeper for high value of the reproduction number and for higher variance of the infectious period. The evolution over time for both the forward and the backward generation intervals show a similar pattern for the different infectious period distributions though we notice that a higher variability is observed for scenarios in which the infectious period distribution has larger variance.

**Fig 1.**
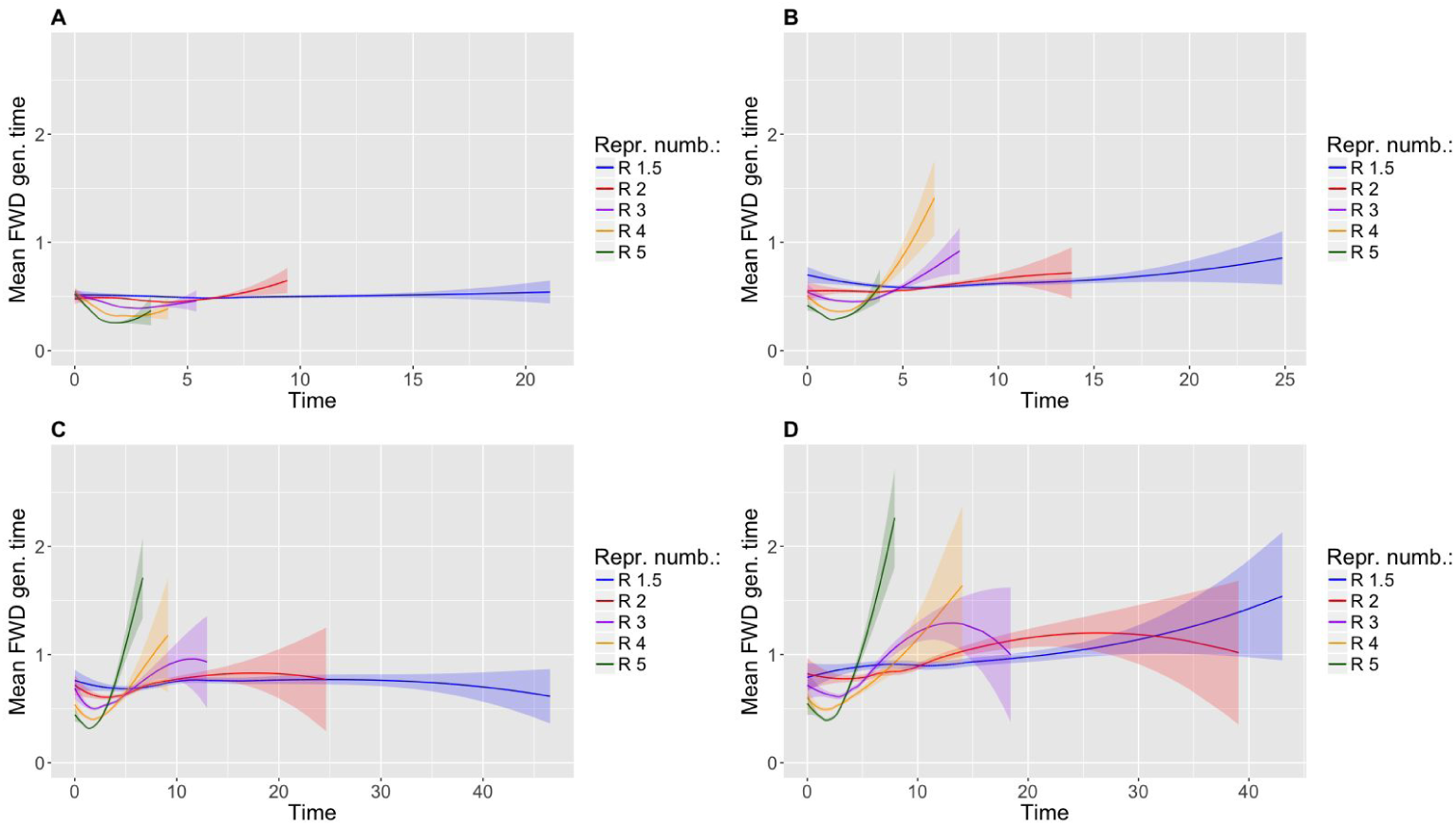
Evolution of the mean forward generation time. (A) Assumes a constant infectious period of unitary length, (B) a gamma infectious period Γ(2, 0.5), (C) an exponential infectious period of rate 1 and (D) a gamma infectious period Γ(0.5, 2).

**Fig 2.**
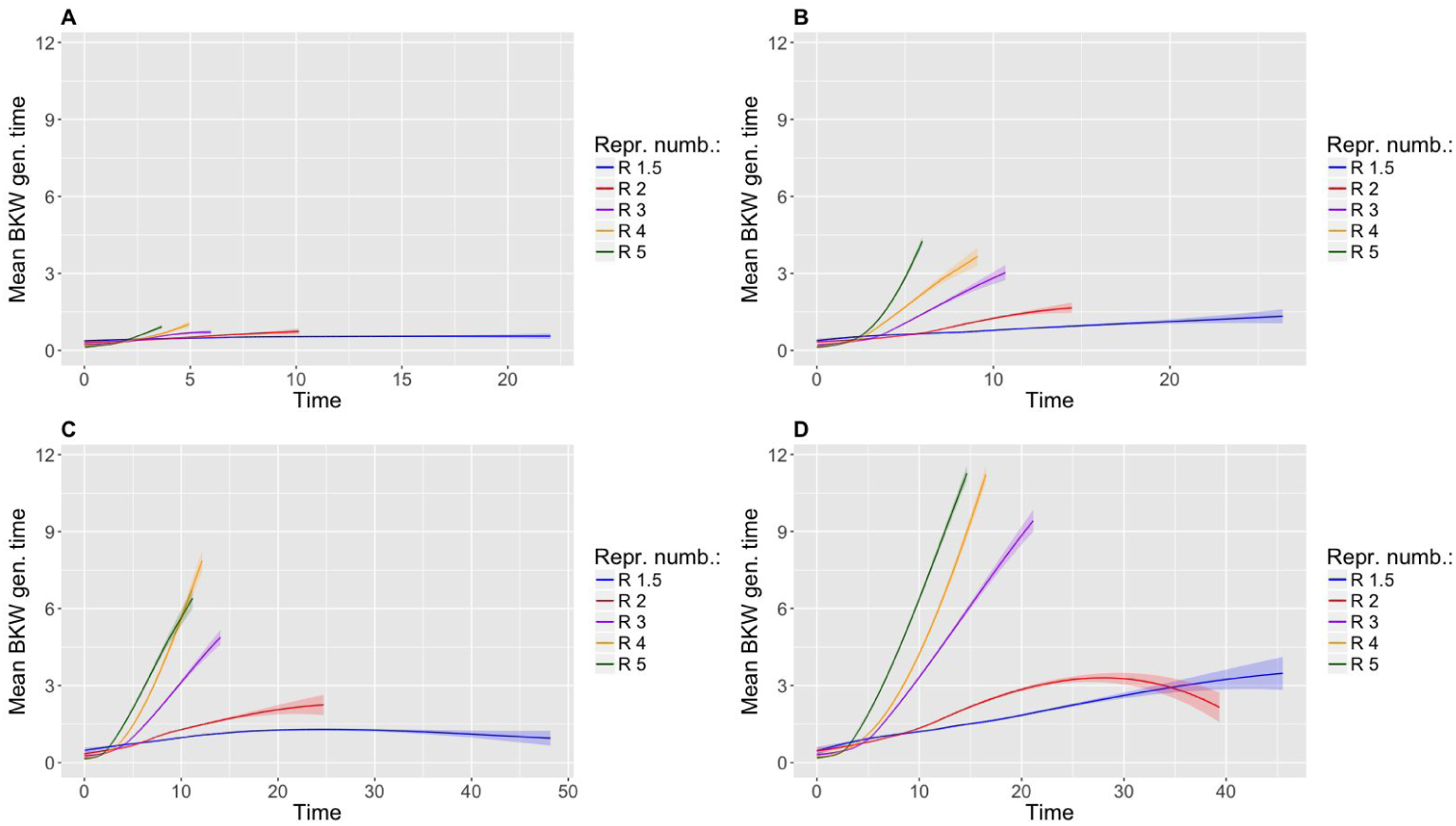
Evolution of the mean backward generation time. (A) Assumes a constant infectious period of unitary length, (B) a gamma infectious period Γ(2, 0.5), (C) an exponential infectious period of rate 1 and (D) a gamma infectious period Γ(0.5, 2).

Lastly, we observe that the different population sizes do not affect considerably the average value of the forward and backward generation time (Table 1). In Fig S5 we show that also the evolution over time is similar for the different sizes considered in the paper.

### Contraction of the mean forward generation time

In this section, our focus is on the evolution over time of the mean forward generation time and on the impact of competition and depletion thereon. We firstly show that competition and depletion are present in our model reporting, respectively, the mean value of the generations where competition is present and the variation in the number of susceptible individuals.

Results reported in Table 2 show that when the reproduction number increases, also the number of generations affected by competition and the mean number of competitors increase. Furthermore, *p*_*c*_ is stable for the tested population size, while the depletion intensity is more accentuated in small population.

**Table 2.** Competition and depletion intensity

| Pop. size | N = 4 |  |  | N = 10 |  |  | N = 20 |  |  | N = 50 |  |  | N = 100 |  |  | N = 200 |  |  |
| --- | --- | --- | --- | --- | --- | --- | --- | --- | --- | --- | --- | --- | --- | --- | --- | --- | --- | --- |
| $\mathcal{R}_0$ | 1.5 | 3 | 5 | 1.5 | 3 | 5 | 1.5 | 3 | 5 | 1.5 | 3 | 5 | 1.5 | 3 | 5 | 1.5 | 3 | 5 |
| $p_c$ | 0.08 | 0.18 | 0.25 | 0.06 | 0.21 | 0.30 | 0.06 | 0.22 | 0.32 | 0.05 | 0.22 | 0.33 | 0.05 | 0.22 | 0.33 | 0.04 | 0.22 | 0.33 |
| $\mu_c$ | 2.03 | 2.08 | 2.10 | 2.05 | 2.18 | 2.28 | 2.06 | 2.16 | 2.27 | 2.07 | 2.16 | 2.28 | 2.05 | 2.16 | 2.29 | 2.03 | 2.16 | 2.29 |
| $\varphi$ | 3.11 | 3.70 | 5.12 | 0.89 | 1.18 | 1.94 | 0.45 | 0.88 | 1.53 | 0.25 | 0.74 | 1.33 | 0.19 | 0.71 | 1.29 | 0.16 | 0.68 | 1.26 |

Average values of the proportion of generations affected by competition ( $p_c$ ), the mean number of competitors ( $\mu_c$ ) and the depletion intensity ( $\varphi$ ).

In Fig 3 we show the phenomena of competition and depletion over time in a population of size *N* = 500. Each point reported in the graph represents the start of the infectious period of an individual for which one (G-1), two(G-2), three(G-3) or four (G-4) generations are affected by competition. Generations affected by competition are concentrated in the declining phase of the forward generation time, close to the maximum contraction point. In the graph we report also the probability of encountering a susceptible individual and the time at which the depletion is at its maximum; the latter is in the region where contraction is at maximum.

**Fig 3.**
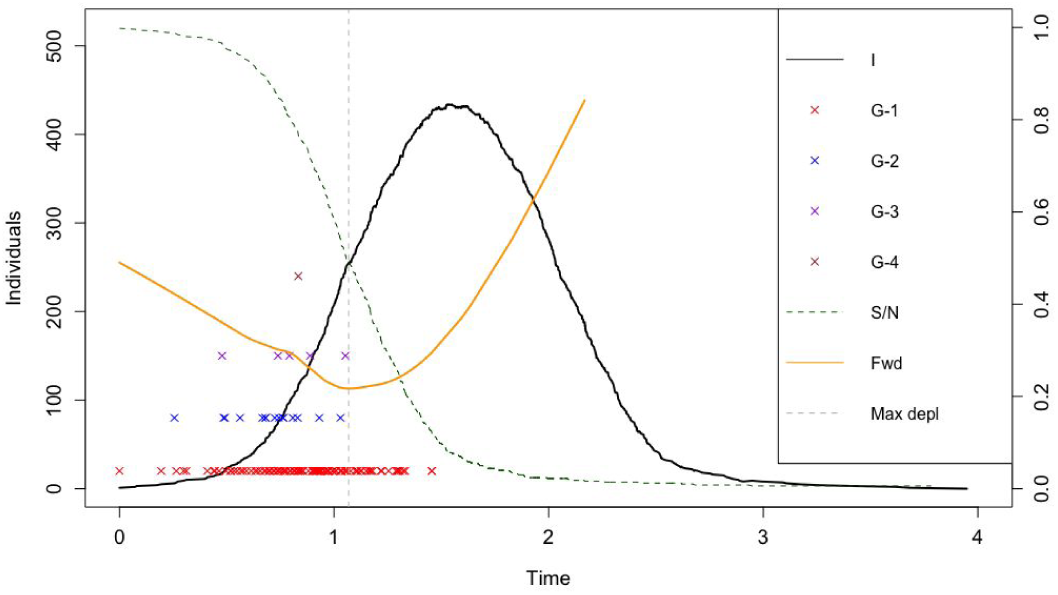
Competition and depletion. Number of infectious individuals over time (black line), mean forward generation time (orange line), proportion of susceptible individuals (green dashed line), time of maximum depletion (grey dashed line) and number of generations affected by competition for a single individual (G-1,G-2,G-3,G-4).

In the vaccination scenario individuals are vaccinated at a precise time during the epidemic. In Table 3 we report the values of the epidemic characteristics for different vaccination times in a population of size *N* = 1000 and a reproduction number of 1.5. When the vaccination coverage is high, i.e. 90%, the mean values are always smaller with respect to the baseline scenario, independently from the vaccination time. This difference is higher when vaccination takes place in the increasing phase of the epidemic (*t*_*v*_ = 3) and in the main phase (*t*_*v*_ = 5). Vaccination in the last part does not affect remarkably the mean values of the generation time distributions. When the vaccination is performed in the early stage of an epidemic (*t*_*v*_ = 2), the impact is clearly visible for high vaccination coverage (60%, 90%) while for a small coverage the impact seems negligible.

**Table 3.** Epidemic characteristics - vaccination scenario

| Vacc. time | $t_v = 2$ | | | $t_v = 3$ | | | $t_v = 5$ | | | $t_v = 7$ | | | $t_v = 9$ | | |
| --- | --- | --- | --- | --- | --- | --- | --- | --- | --- | --- | --- | --- | --- | --- | --- |
| Coverage | 30% | 60% | 90% | 30% | 60% | 90% | 30% | 60% | 90% | 30% | 60% | 90% | 30% | 60% | 90% |
| $T_{max}$ | 12.46 | 5.92 | 3.84 | 11.75 | 6.70 | 4.83 | 11.38 | 8.27 | 6.79 | 11.53 | 9.75 | 8.66 | 11.89 | 11.04 | 10.35 |
| $FS$ | 177.57 | 120.55 | 113.85 | 211.02 | 146.52 | 136.48 | 329.86 | 271.18 | 245.43 | 452.72 | 420 | 394.23 | 539.33 | 516.17 | 508.79 |
| $\omega_f$ | 0.49 | 0.44 | 0.38 | 0.49 | 0.45 | 0.41 | 0.48 | 0.47 | 0.44 | 0.49 | 0.48 | 0.47 | 0.49 | 0.48 | 0.48 |
| $\omega_b$ | 0.49 | 0.45 | 0.40 | 0.49 | 0.46 | 0.43 | 0.49 | 0.47 | 0.45 | 0.49 | 0.48 | 0.47 | 0.49 | 0.48 | 0.48 |
| $p_c$ | 0.01 | 0.02 | 0.03 | 0.04 | 0.02 | 0.02 | 0.03 | 0.03 | 0.04 | 0.04 | 0.04 | 0.04 | 0.04 | 0.04 | 0.04 |
| $\mu_c$ | 2.01 | 2 | 1.9 | 2.02 | 2.01 | 2.01 | 2.03 | 2.02 | 2.02 | 2.02 | 2.02 | 2.02 | 2.02 | 2.03 | 2.03 |

Average values of duration ( $T_{max}$ ), final size ( $FS$ ), mean forward ( $\omega_f$ ) and mean backward ( $\omega_b$ ) generation times together with the proportion of generations affected by competition ( $p_c$ ), mean number of competitors ( $\mu_c$ ) and depletion intensity ( $\varphi$ ).

In Fig 4, we report the simulated evolution over time of the mean forward generation interval for the vaccination times *t*_*v*_ = 5. We can appreciate the phenomenon of contraction that increases with an increasing in the vaccination coverage reaching the maximum contraction right before the vaccination time. In S1 Appendix we report the forward generation time in case of *t*_*v*_ = 2, 9 showing a similar trend to the plot reported hereunder (S2 Fig).

**Fig 4.**
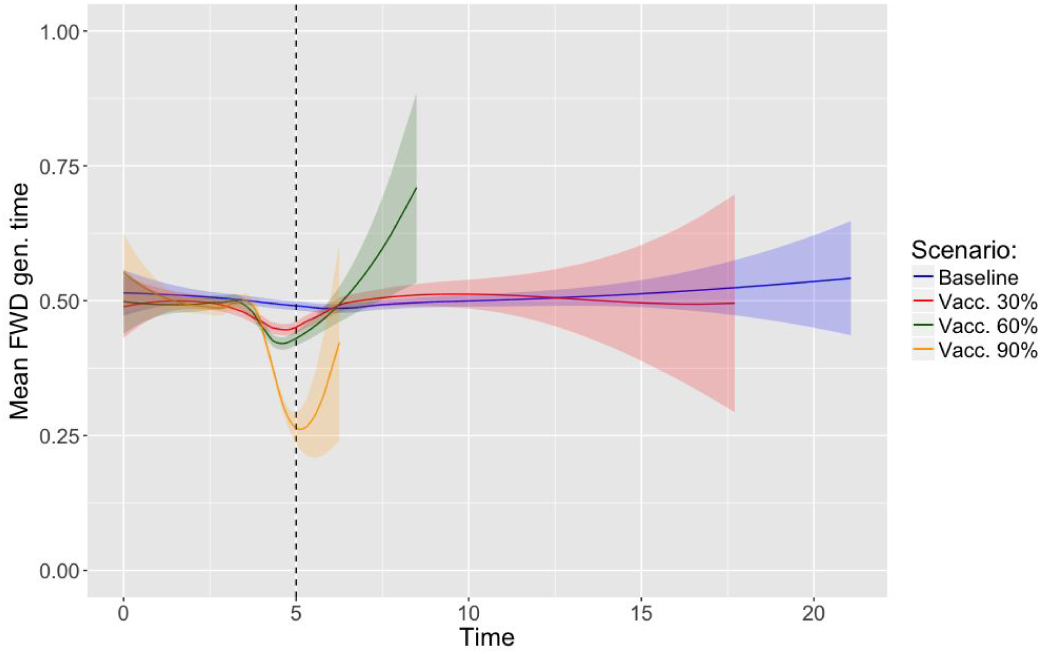
Evolution of the mean forward generation time - Vaccination scenario. Comparison of the evolution of the mean forward generation time between the baseline scenario (blue line) and the vaccination scenarios. The vaccination time is set at *t*_*v*_ = 5 and the vaccination coverages considered are 30% (red line), 60% (green line) and 90% (orange line).

In the pure depletion scenario the phenomenon of competition is not present anymore. We compare Table 4 (pure depletion scenario) with Tables 1 and 5 (baseline scenario) to assess the impact of competition. Results show that there are no remarkable differences among these two scenarios, neither for different values of the reproduction number nor for different population sizes. We observe that the mean values of the generation time distributions are slightly smaller when competition is present, both in the forward and in the backward scheme.

**Table 4.** Epidemic characteristics - Pure depletion scenario

| Pop. size | N = 100 |  |  |  |  | N = 500 |  |  |  |  | N = 1000 |  |  |  |  |
| --- | --- | --- | --- | --- | --- | --- | --- | --- | --- | --- | --- | --- | --- | --- | --- |
| $R_0$ | 1.5 | 2 | 3 | 4 | 5 | 1.5 | 2 | 3 | 4 | 5 | 1.5 | 2 | 3 | 4 | 5 |
| $T_{max}$ | 7.06 | 5.75 | 3.89 | 3.11 | 2.73 | 11.26 | 7.59 | 4.78 | 3.72 | 3.23 | 12.70 | 8.35 | 5.19 | 3.97 | 3.42 |
| $FS$ | 61.56 | 86.96 | 98.93 | 99.96 | 100 | 315.60 | 434.84 | 494.77 | 499.86 | 500 | 629.80 | 868.91 | 989.73 | 999.73 | 999.99 |
| $\omega_f$ | 0.49 | 0.47 | 0.41 | 0.35 | 0.29 | 0.49 | 0.47 | 0.42 | 0.35 | 0.30 | 0.49 | 0.48 | 0.42 | 0.35 | 0.30 |
| $\omega_b$ | 0.49 | 0.47 | 0.42 | 0.36 | 0.31 | 0.49 | 0.48 | 0.42 | 0.36 | 0.31 | 0.49 | 0.48 | 0.42 | 0.36 | 0.31 |
| Max loess | 0.68 | 0.65 | 0.61 | 0.56 | 0.51 | 0.63 | 0.62 | 0.60 | 0.56 | 0.54 | 0.61 | 0.61 | 0.60 | 0.55 | 0.54 |
| Min loess | 0.33 | 0.33 | 0.27 | 0.20 | 0.16 | 0.39 | 0.38 | 0.31 | 0.22 | 0.17 | 0.40 | 0.40 | 0.33 | 0.23 | 0.18 |
| Range loess | 0.34 | 0.32 | 0.34 | 0.36 | 0.35 | 0.23 | 0.25 | 0.28 | 0.34 | 0.37 | 0.20 | 0.22 | 0.27 | 0.32 | 0.36 |
| $\varphi$ | 0.21 | 0.43 | 0.85 | 1.20 | 1.52 | 0.16 | 0.39 | 0.83 | 1.17 | 1.48 | 0.16 | 0.40 | 0.82 | 1.16 | 1.48 |

Average values of duration ( $T_{max}$ ), final size ( $FS$ ), mean forward ( $\omega_f$ ) and mean backward ( $\omega_b$ ) generation times, maximum, minimum and range of the loess regression (Max loess, Min loess and Range loess) and depletion intensity ( $\varphi$ ).

**Table 5.** Loess regression values and depletion intensity - Baseline scenario

| Pop. size | N = 100 |  |  |  |  | N = 500 |  |  |  |  | N = 1000 |  |  |  |  |
| --- | --- | --- | --- | --- | --- | --- | --- | --- | --- | --- | --- | --- | --- | --- | --- |
| $\mathcal{R}_0$ | 1.5 | 2 | 3 | 4 | 5 | 1.5 | 2 | 3 | 4 | 5 | 1.5 | 2 | 3 | 4 | 5 |
| Max loess | 0.69 | 0.66 | 0.62 | 0.58 | 0.53 | 0.62 | 0.63 | 0.63 | 0.60 | 0.56 | 0.60 | 0.61 | 0.62 | 0.61 | 0.58 |
| Min loess | 0.33 | 0.31 | 0.27 | 0.21 | 0.17 | 0.39 | 0.38 | 0.32 | 0.26 | 0.21 | 0.41 | 0.39 | 0.34 | 0.28 | 0.22 |
| Range loess | 0.36 | 0.35 | 0.35 | 0.37 | 0.37 | 0.23 | 0.25 | 0.30 | 0.33 | 0.35 | 0.19 | 0.22 | 0.28 | 0.33 | 0.36 |
| $\varphi$ | 0.19 | 0.36 | 0.71 | 1.02 | 1.29 | 0.14 | 0.32 | 0.67 | 0.97 | 1.25 | 0.13 | 0.32 | 0.69 | 0.97 | 1.24 |

Average values of maximum, minimum and range of the loess regression (Max loess, Min loess and Range loess) and depletion intensity ( $\varphi$ ).

In Fig 5 we report the mean forward generation interval over time comparing the pure depletion and the baseline scenarios in the case of *ℛ*_0_ = 5, where the number of generations affected by competition is higher. However, the evolution of the curves result to be similar, showing a little difference between the two considered scenarios.

**Fig 5.**
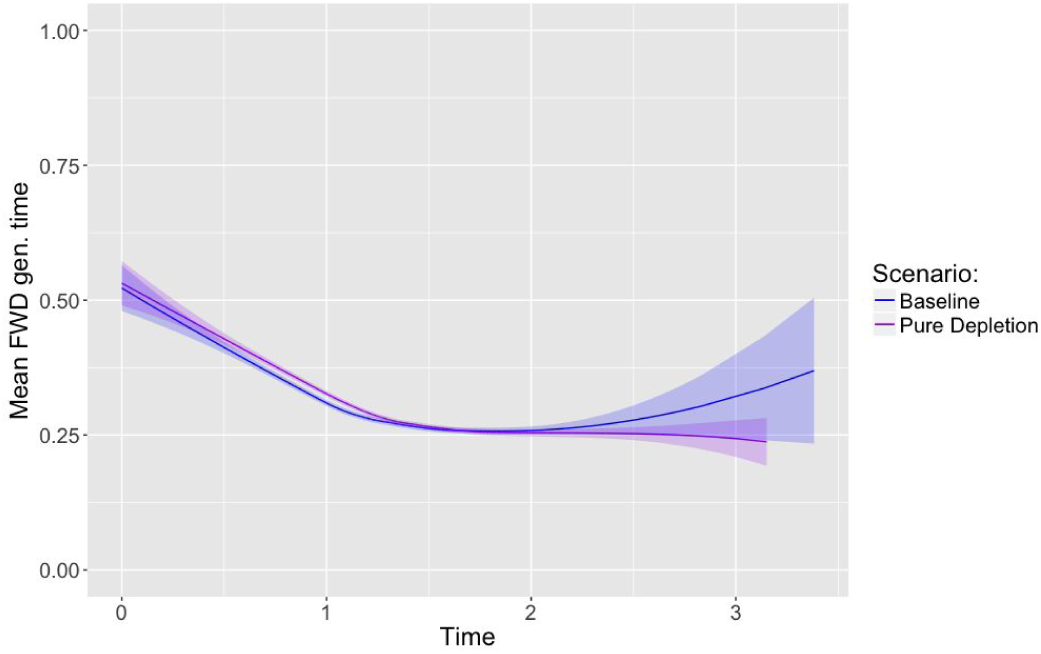
Evolution of the mean forward generation time - Pure depletion scenario. Comparison of the evolution of the mean forward generation time between the baseline scenario (blue line) and the pure depletion scenarios (purple line).

In the competition scenario, we increase competition in the time interval (3, 7), for a population of size *N* = 1000 and a reproduction number of value *ℛ*_0_ = 1.5. In Table 6 we report the epidemic characteristics; we notice that the mean value of the generation time is smaller compared to the one in the baseline scenario (Table 1), both for the forward and the backward scheme. We also notice that the mean number of generation affected by competition and the mean number of competitors is higher respect to the baseline scenario with same reproduction number (Table 2).

**Table 6.**
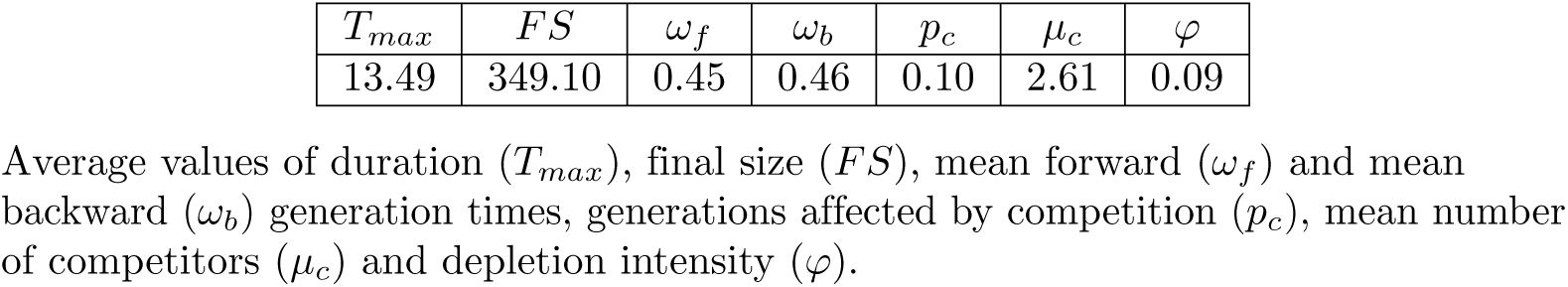
Epidemic characteristics - Competition scenario

| $T_{max}$ | $FS$ | $\omega_f$ | $\omega_b$ | $p_c$ | $\mu_c$ | $\varphi$ |
| --- | --- | --- | --- | --- | --- | --- |
| 13.49 | 349.10 | 0.45 | 0.46 | 0.10 | 2.61 | 0.09 |

Average values of duration ( $T_{max}$ ), final size ( $FS$ ), mean forward ( $\omega_f$ ) and mean backward ( $\omega_b$ ) generation times, generations affected by competition ( $p_c$ ), mean number of competitors ( $\mu_c$ ) and depletion intensity ( $\varphi$ ).

In Fig 6 we show the evolution of the forward generation interval comparing the competition and the baseline case. It is clearly visible that the mean forward generation time contracts around the interval in which the competition intensity is increased.

**Fig 6.**
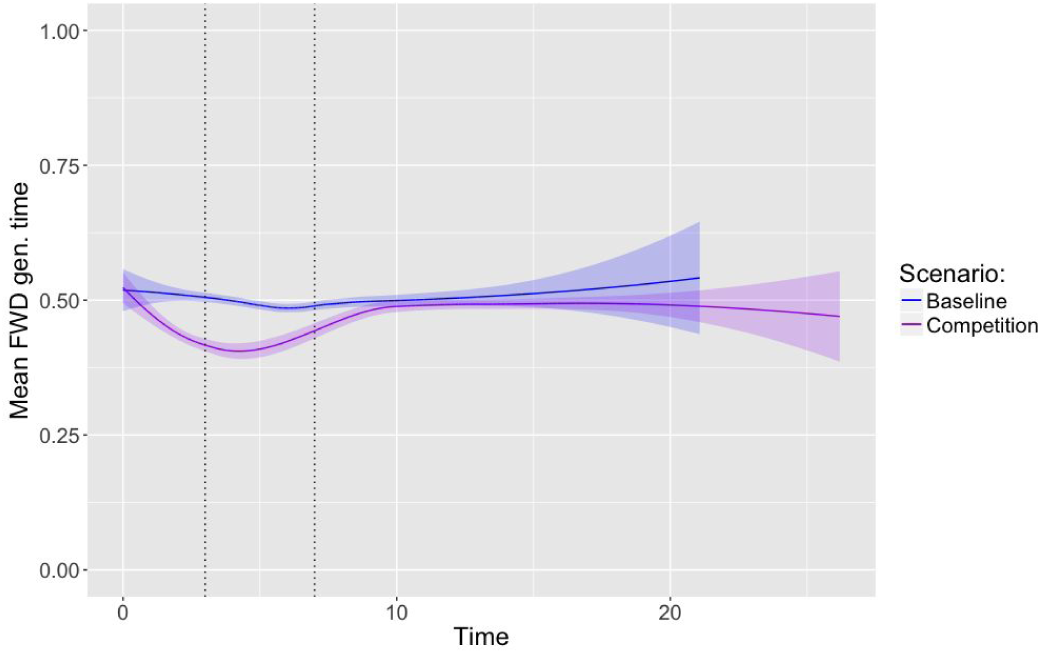
Evolution of the mean forward generation time - Competition scenario. Comparison of the evolution of the mean forward generation time between the baseline scenario (blue line) and the competition scenario (purple line). The dotted lines represent the interval in which the competition is increased.

## Discussion

The present study was designed with the aim of characterizing the main cause of contraction in the evolution over time of the mean forward generation interval. The results of this investigation address both depletion of susceptible individuals and competition among infectors as causes of contraction. Scenarios where the depletion or the competition effects are increased show that the mean forward generation interval contracts with a peak close to the time point of maximum depletion or maximum competition, even for a small value of the reproduction number. In the pure depletion scenario, depletion clearly causes the phenomenon of contraction. Given that results are similar to the baseline scenario where depletion and competition are both present, we can conclude that depletion is predominant in shaping the evolution over time of the mean forward generation time.

A rapid depletion forces the infectious person to make more shorter infectious contacts because the probability of encountering a susceptible individual drastically decreases over time. This phenomenon is highlighted for high values of the reproduction number: the rate of the contact process is higher implying that infectious individuals propose on average more infectious contacts during their infectious period. The higher depletion effect increases the proportion of short contact intervals resulting in a faster and larger contraction of the mean forward generation time.

Competition also affects the generation time distribution but its effect is directed to a single generation. To have a similar impact on the mean forward generation time as caused by depletion, competition should affect most of all the generations that an individual makes. This is simulated in the scenario where competition is increased and results show a potentially large impact of competition on the forward generation time. However, in a baseline scenario where competition is not increased rarely more than one generation per single individual is affected by competition and the mean number of competitors is not particularly high to be able to explain a decreasing mean forward generation time (Fig 3). Furthermore, the effect of competition is strongly dependent on the competitor’s infection time and does not not always affect the considered generation. The contact interval that leads to the generation time can be the longest one among the contact intervals proposed by the infectors since the next generation is the minimum of the set given by the infection times plus the proposed contact times. Competition is slightly affecting the forward and backward generation time: Table 6 shows a small decrease probably due to the competition effect.

This paper’s focus is on the evolution over time of the mean forward generation time but the backward generation scheme is of interest too. The mean backward generation time is known to be increasing [6, 20] and differently from the forward scheme a single generation is considered for every time point. The increasing trend is due to the fact that the probability of encountering a susceptible decreases over time, but a more intense competition can, also in this case, modify the evolution of the mean value over time. This is shown in Fig S6, where in a preliminary investigation the baseline, vaccination, increasing competition and pure depletion scenarios are compared. The effect of competition clearly flattens the increasing trend of the mean backward generation time over time while increased depletion (vaccination scenario) causes a steeper increase.

Another aim of this study was to investigate the effect of the reproduction number, the infectious period and the population size on the mean generation time in the backward and forward scheme. Results show that their mean values over time decrease faster and more with increasing reproduction number. Furthermore, the infectious period distribution affects remarkably these mean values with a higher impact when the variance of the selected infectious period distribution is higher. This finding can be explained with the mathematical framework developed by Nishiura (2010) where he mathematically relates the probability density function of the generation interval to the infectious period distribution. The population size does not remarkably influence the mean generation intervals for the tested sizes.

Although we have looked at compartmental SIR models, we expect our conclusion to hold for more complicated compartmental models, and even for epidemics models on structured contact networks. A limitation of our investigation is the assumption made for the infectious contact process to be described by a Poisson process and to be homogeneous in the population: in a structured population the infectious contact process depends on the location where the contacts take place because of different behaviour of individuals yielding different contact processes [21].

The findings of the present study clearly show the non-constant behaviour of both backward and forward generation interval, in line with the literature [5, 6, 8, 20]. Moreover, this has been the first attempt to thoroughly examine the cause of the contraction: competition and depletion are both capable of affecting the evolution over time of the mean generation interval. As result, in such models, estimators of the generation time distribution, accounting for both depletion of susceptible and competing-risk, are to be preferred.

## Supporting information

### Stepwise algorithm

Consider an introductory case in an entirely-susceptible population of size *N* at time *t*_(1)_ = 0 and assume the person recovery period is known. The epidemic evolves in the following way: the introductory case makes contacts with all the susceptible person in the population {*τ*_(1)_*j*__ : *j* ∈ *S*_*t*_(1)__} according to a contact interval distribution with hazard function *h*_(1)_*j*__ (*τ*). Among all contacts made by the introductory case, only the infectious contact have potential to generate the secondary cases. Set 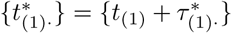 as the proposed infection times of all the infectious contacts made by the first case, where 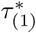. denotes all the infectious contact intervals of the first case. Note that all recipients of these infectious contacts will be infected at or before time 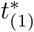, either from person (1) or from another infector. In fact, the second infected case corresponds to the smallest proposed infection time and occurs at time 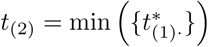. Now there are two infected persons in the population. Similarly, the second case makes contacts {*τ*_(2)_*j*__ : *j ∈ S*_*t*_(2)__} with the remaining *N* − 2 susceptible persons according to the hazard function *h*_(2)_*j*__ (*τ*). Set 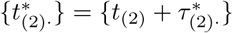 as the proposed infection times. The third case occurs at the minimum proposed infection time between the available infectious contacts made by the first and second infected case: 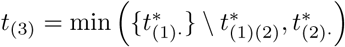. Note that all proposed infectious time for already-infected cases has to be omitted. The third case can be infected either by the first or the second infected case, depending on which one has made the first infectious contact. The epidemic continues until there are no infectious persons. We summarize the algorithm in 5 steps.

For the *i*th infected person:

1. Generate contact intervals {*τ*_(*i*)*j*_: *j ∈ S*_*t*_(*i*)__} according to the hazard function *h*_(*i*)*j*_(*τ*).
2. Record the proposed infection time 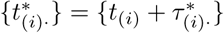.
3. Recursively, set 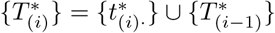, where 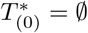.
4. The next infected case occurs at time 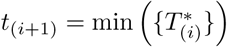.
5. Set 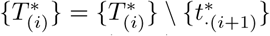, where 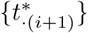 is the set of all proposed infection times for the (*i* + 1)-th case from all the potential infectors.

The outbreak ends when there are no longer infectious persons. This algorithm highlight the phenomenon of competition among infectious persons (Kenah et al. 2008).

### Parallel algorithm

The other algorithm for generating epidemics is the parallel algorithm by Scalia Tomba et al. (2010). The epidemic begins with an imported infection from outside the population at time *t*_(1)_ = 0. The introductory case has an infectious period of length *ℛ*_(1)_, according to a desirable statistical distribution *F*. We assume that the total number of contacts for a fixed length of the infectious period *ℛ*, follows a homogeneous Poisson process with constant intensity *βr*, where *β* is a known constant. Consequently, the inter-arrival contact times for an infected case *i*, denoted by {*δ*_*i*1_, *δ*_*i*2_, …} are independent and follow an exponential distribution of parameter *βr*. The introductory case will randomly meet the first individual at time 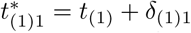. If this contact happens within the infectious period *ℛ*_(1)_, i.e. 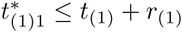, it is an infectious contact and the infection time of the next case is 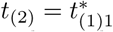. At this stage, the newly infected case and its infector will contact other people randomly in the population. Unlike the stepwise algorithm, this contact can be potentially generated also for already infected person. In parallel, the first case will generate his or her second contact at time 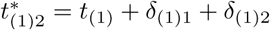 and the second case will generate his or her contact at time 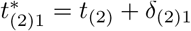. As long as the contacts are made with susceptible persons and lie within the respective infectious period, they will be infectious contact. The third case will correspond to an infectious contact with the earliest contact time. The epidemic grows in this way until there are no infectious person anymore. We summarize the parallel algorithm:

For each infectious case *i*:

1. Generate the next inter-arrival contact time *δ*_*i*+_
2. Set 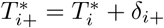 as the proposed next contact time of person *i* where 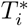 is the previous contact time made by case *i*.
3. If 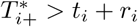 the case *i* is already recovered and the contact can not be an infectious contact, then discard it. If the contact is made with a susceptible person it is a proposed infectious time and record 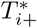.
4. The next infectious time is the minimum over all the proposed infectious time.
5. Generate the recovery time for the newly infected.

### Comparison of the simulation algorithms

The two algorithms give the same results when looking at the mean value of the considered epidemic characteristics and when plotting the evolution over time of the mean forward and backward generation times. In S1 Table we reported the average quantities of mean duration *T*_*max*_, mean final size *FS*, mean forward generation time 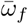 and mean backward generation generation time 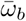 for simulations with the stepwise algorithm. Values are almost the same as the ones reported for the parallel case in Table 1. In S1 Fig we compare the evolution over time of the mean forward generation time in the vaccination scenario for *ℛ*_0_ = 1.5, 3 varying the infectious period and using both parallel and stepwise algorithms. The plot shows that the results are independent from the selected algorithm and infectious period. Same conclusions hold also for the gamma infectious period distributions considered in this paper.

### Maximum depletion

Every time an infectious contact is proposed we keep track of the current time *t*_*j*_ and the probability of encountering a susceptible individual at that specific time *ξ*_*t*_*j*__, computed as the proportion of susceptibles over the population size. We than fit a 5-parameters logistic curve *f*(*x*|{*t*_*j*_}_*j*_, {*ξ*_*t*_*j*__}*j*) to the simulated values using the function *drm* of the *drc* package in R. At this point we approximate the derivative of the fitted function as:

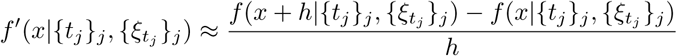

where in our simulation *h* is set to be *h* = 0, 001. According to this discretization, for a specific simulation we define the maximum depletion to be:

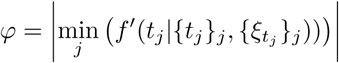

We then compute the mean of the depletion intensity value among all the non extinct simulations.

### Vaccination time

Vaccination affects the phenomenon of contraction differently, depending on the vaccination time. In S2 Fig we report plots for two different vaccination times *t*_*v*_ = 2, 9 in the scenario with *ℛ*_0_ = 1.5 reported in the results section representing the ascending phase and the descending phase of an epidemic outbreak. Even if the intensity is different the phenomenon of contraction is always present.

### Regression of the simulated data

The simulated data are analyzed in R; generations resulting from the first *n* non-extinct simulations are merge together in a unique data file and we predict the value of a loess regression to express the evolution over time of mean forward or backward generation time. This analysis required a huge computational memory, limiting the number of simulations that can be considered. However, this R function allows us to construct a confidence interval to quantify the variability of the loess regression used. In S3 Fig we show how different numbers of considered simulations affect the results. The quality of the fit is based on the number of generations considered and this number is directly related to the population size. We present the evolution over time for the mean forward generation time in a population of size *N* = 1000 for a constant infectious period. These plots enforce the analysis we conducted even if a limited number of simulations are considered.

Another important aspect in the loess regression is the considered span. The fitting is done locally and the considered neighbourhood of a point *x* is based on the value of the span, parameter that has to be given as input in the loess function. In S4 Fig we report the evolution over time of the mean forward generation time for *ℛ* = 1.5 in the vaccination scenario for three different values of the span parameter: 0.5, 0.75(default value) and 1.5 in the case of exponential infectious period. The plot shows differences between the curves in the last part of the outbreak after the vaccination time where few generations are registered. This is probably due to the little number of generations that happen after the vaccination time. We want to remark that the result of contraction holds independently on the type of span considered.

Lastly, we want to show the evolution over time of the mean forward generation time for different population sizes. In S5 Fig we report this quantity for population of size *N* = 100, 500 and 1000 in the case of exponential infectious period. We notice that one plot is the re-scaling of the others; the general evolution of the forward generation intervals is not affect by the population size. The same results apply for the backward generation interval.

### Effect of competition and depletion on the mean backward generation time

In a preliminary study, we investigated also the effect of competition on the mean backward generation time. In S6 Fig we report the evolution over time of the mean backward generation interval *ℛ*_0_ = 1.5, 3, 5 comparing the baseline, the vaccination, the competition and the pure depletion scenario. We note an overall relevant impact of competition when its effect is incremented (competition scenario), registering a smoothed increase. Furthermore, for *ℛ*_0_ = 3, 5 we observe that the pure depletion and the baseline scenario have different evolution indicating an impact of competition.

**S1 Table.**
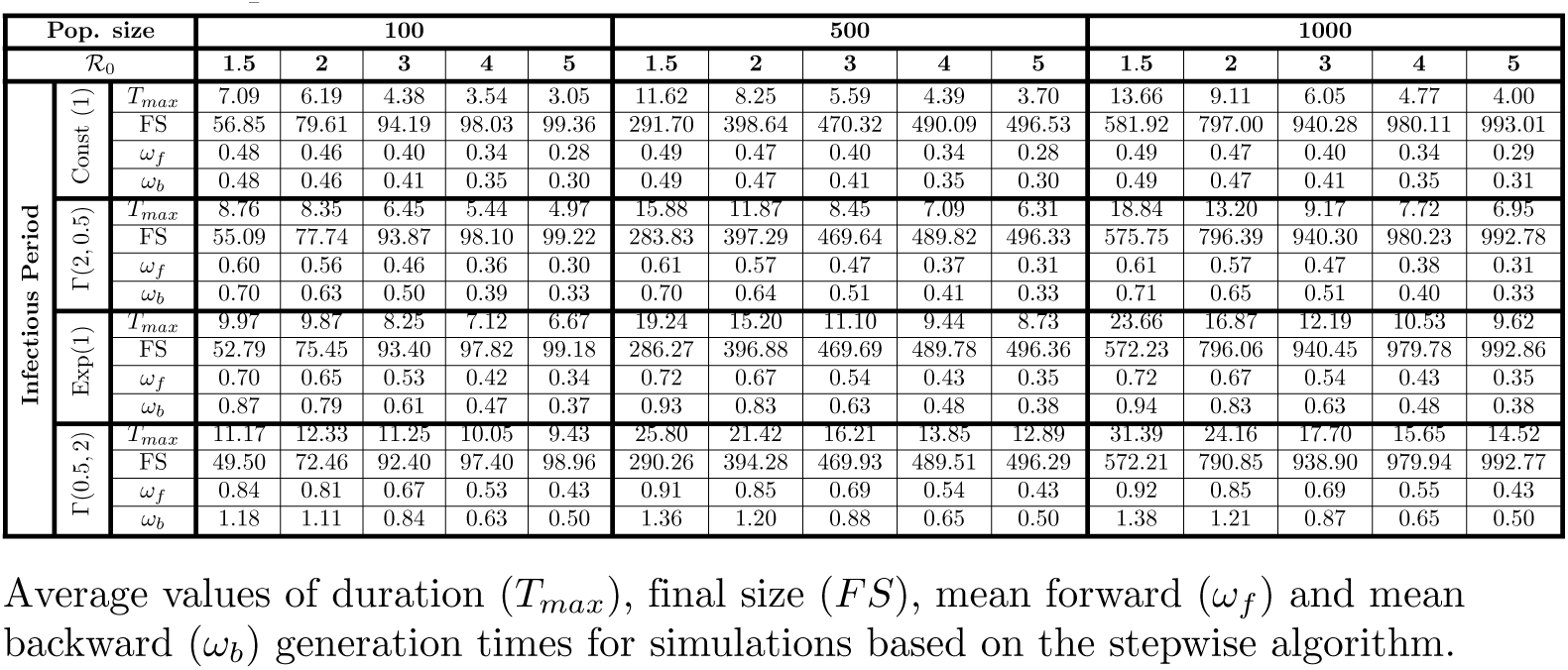
Epidemic characteristics - baseline scenarios

**S1 Fig.**
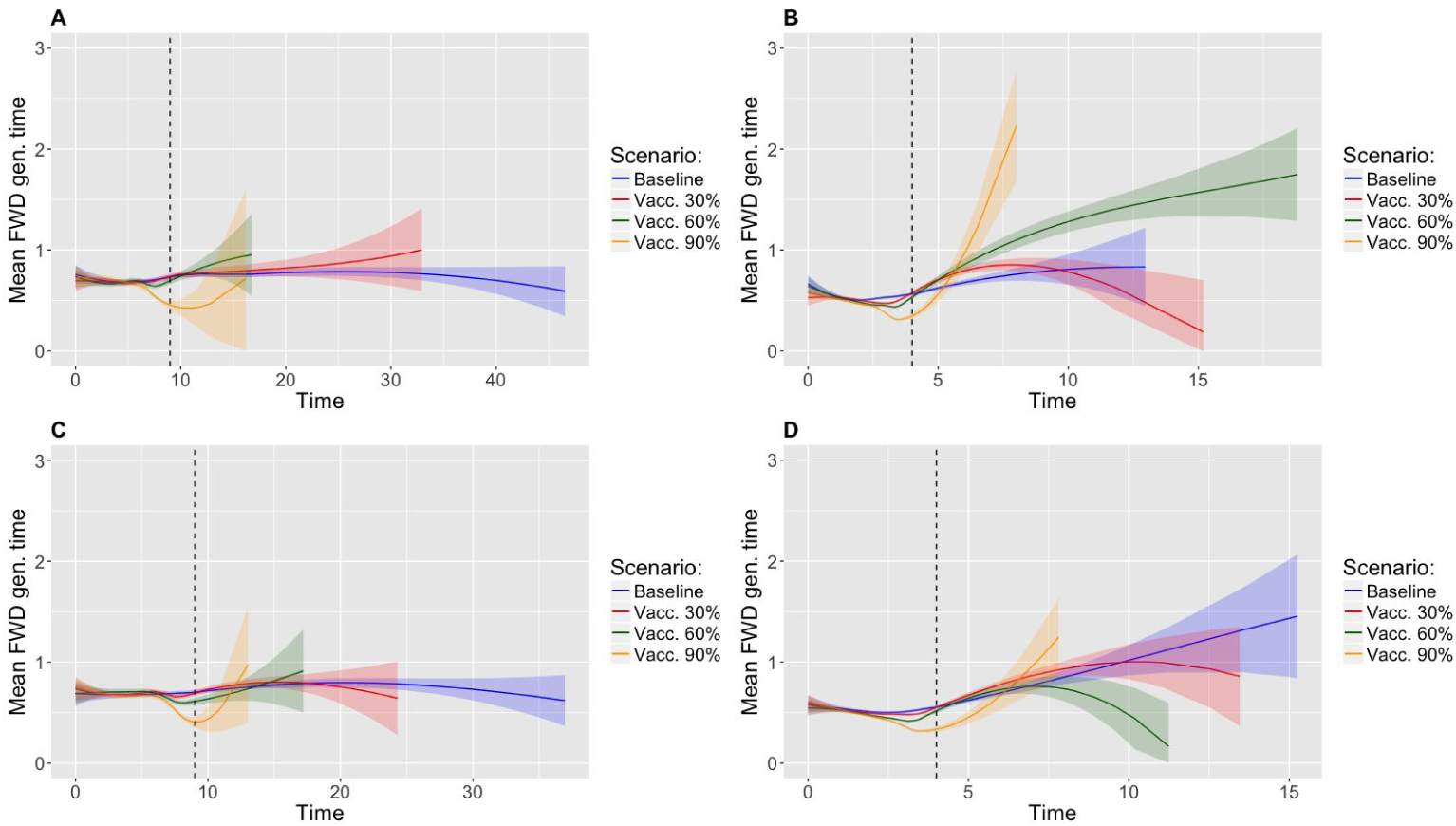
Evolution of the mean forward generation time - Parallel and Stepwise algorithm. Mean forward generation time in the vaccination scenario comparing stepwise and parallel algorithms. (A) and (B) identify the parallel algorithm while (C) and (D) identify the stepwise algorithm. The considered infectious period is exponential with unitary rate and the population is of size *N* = 1000. The reproduction number is of value *ℛ*_0_ = 1.5 for (A) and (C) and of value *ℛ*_0_ = 3 for (B) and (D). The vaccination time is, respectively, *t*_*v*_ = 9 (A and C) and *t*_*v*_ = 5 for (B and D).

**S2 Fig.**
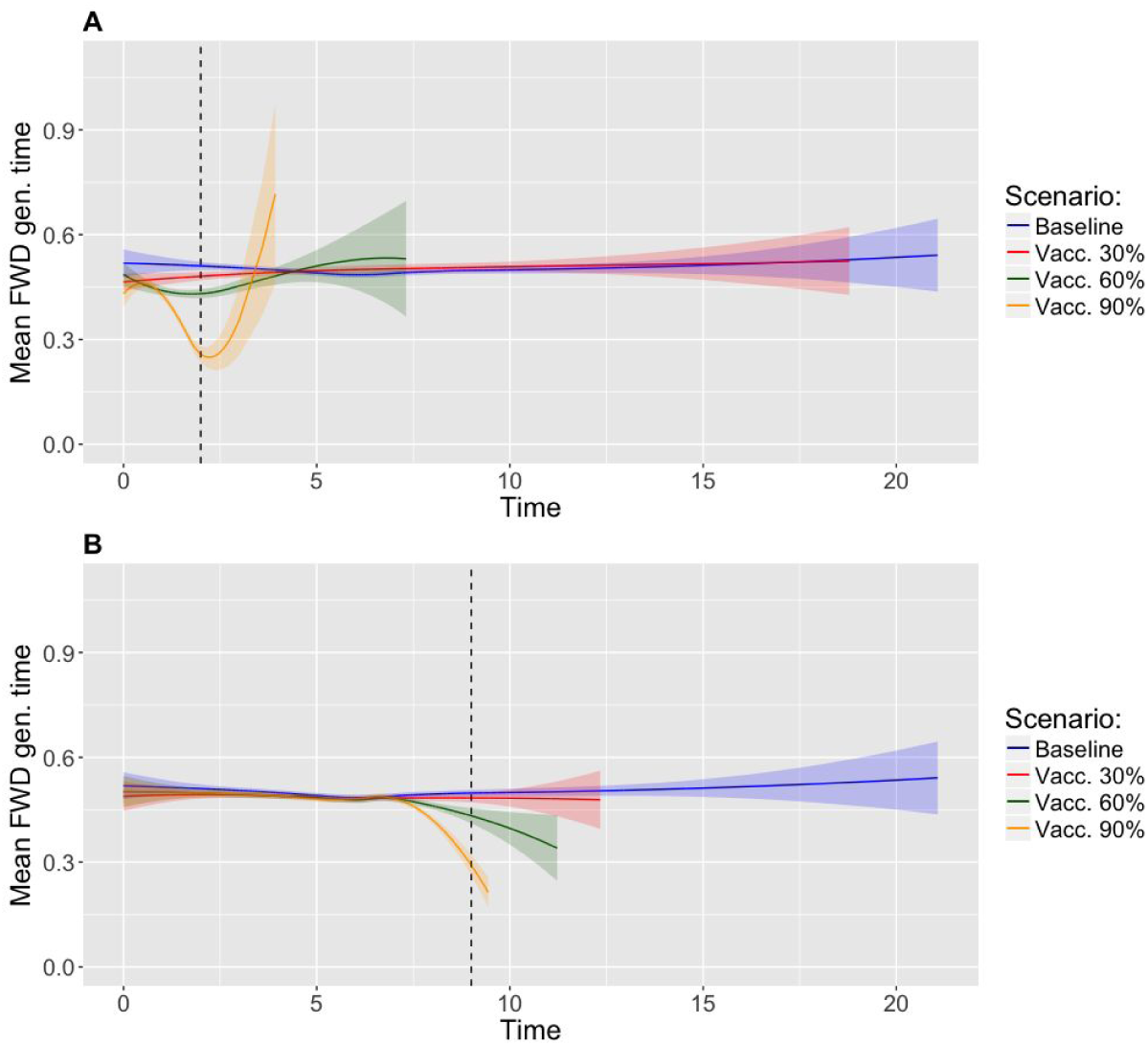
Evolution of the mean forward generation time for different vaccination times. Mean forward generation time for a vaccination time of *t*_*v*_ = 2 (A) and *t*_*v*_ = 9 (B). The population is of size *N* = 1000, the reproduction number of value *ℛ*_0_ = 1.5 and the infectious period is constant.

**S3 Fig.**
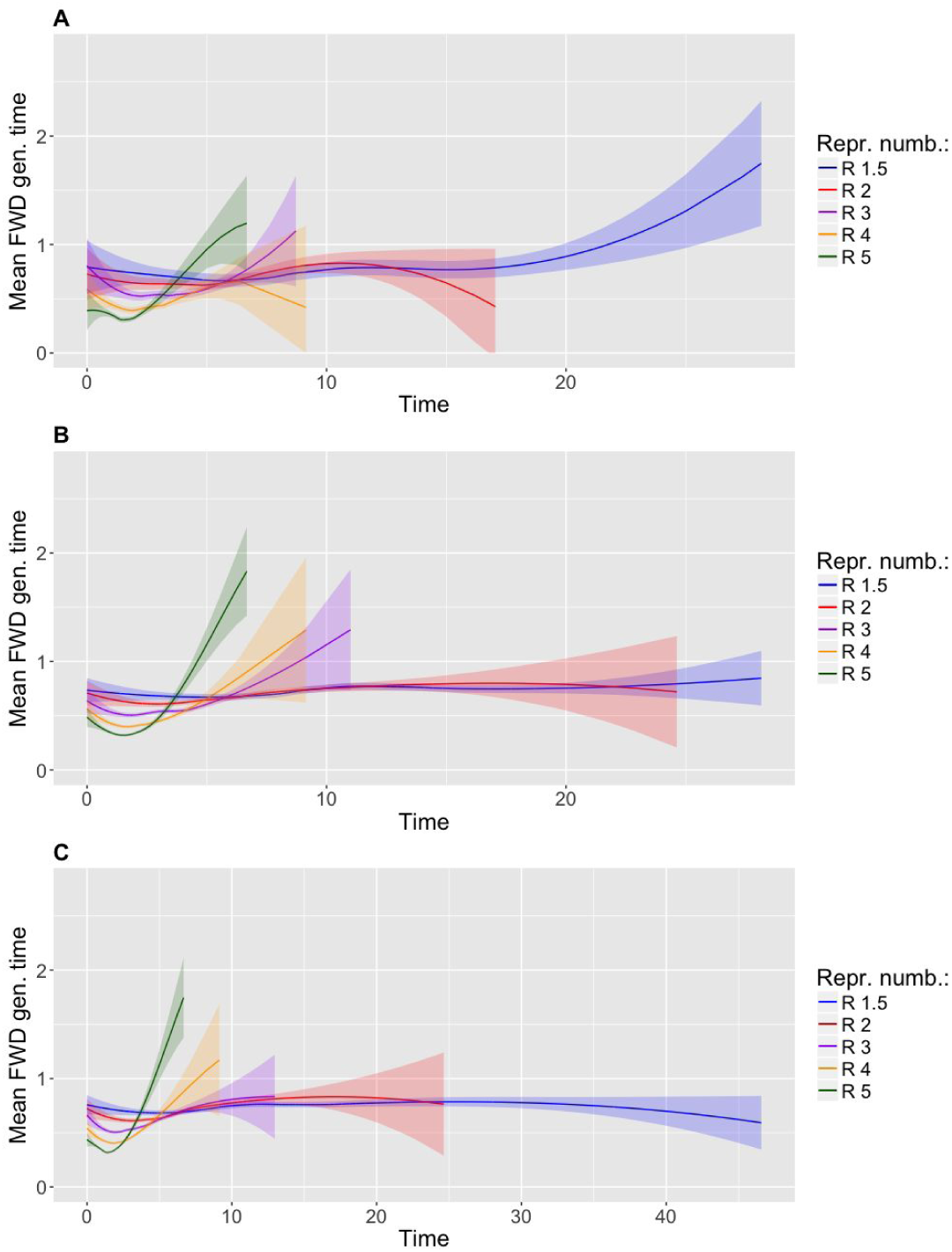
Prediction of the mean forward generation time. Prediction of the mean forward generation time based on, respectively, *n* = 5 (A), *n* = 20 (B) and *n* = 30 non extinct simulations. The selected infectious period is constant with unitary length, the population size is *N* = 1000 and the simulations are obtained with the parallel algorithm. The reproduction number *ℛ*_0_ = 1.5, 2, 3, 4, 5.

**S4 Fig.**
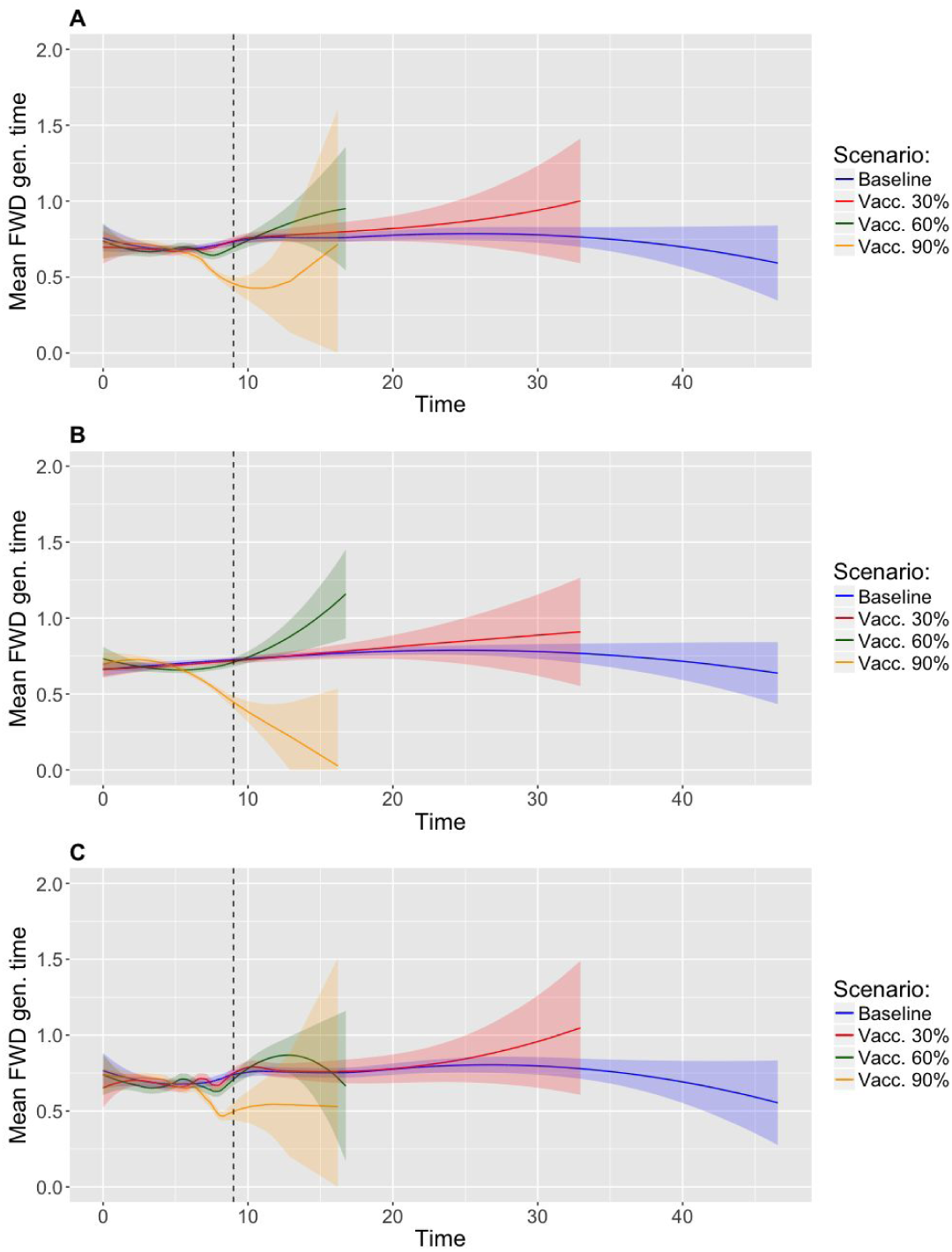
Prediction of the mean forward generation time - varying the span parameter. Prediction of the mean forward generation time based on different value of the span parameter (loess regression). Respectively, span=0.75 (A), span=1 (B) and span=0.5 (C). The population is of size *N* = 1000, the vaccination time is *t*_*v*_ = 9, the reproduction number *ℛ*_0_ = 1.5 and the infectious period is exponential of unitary rate.

**S5 Fig.**
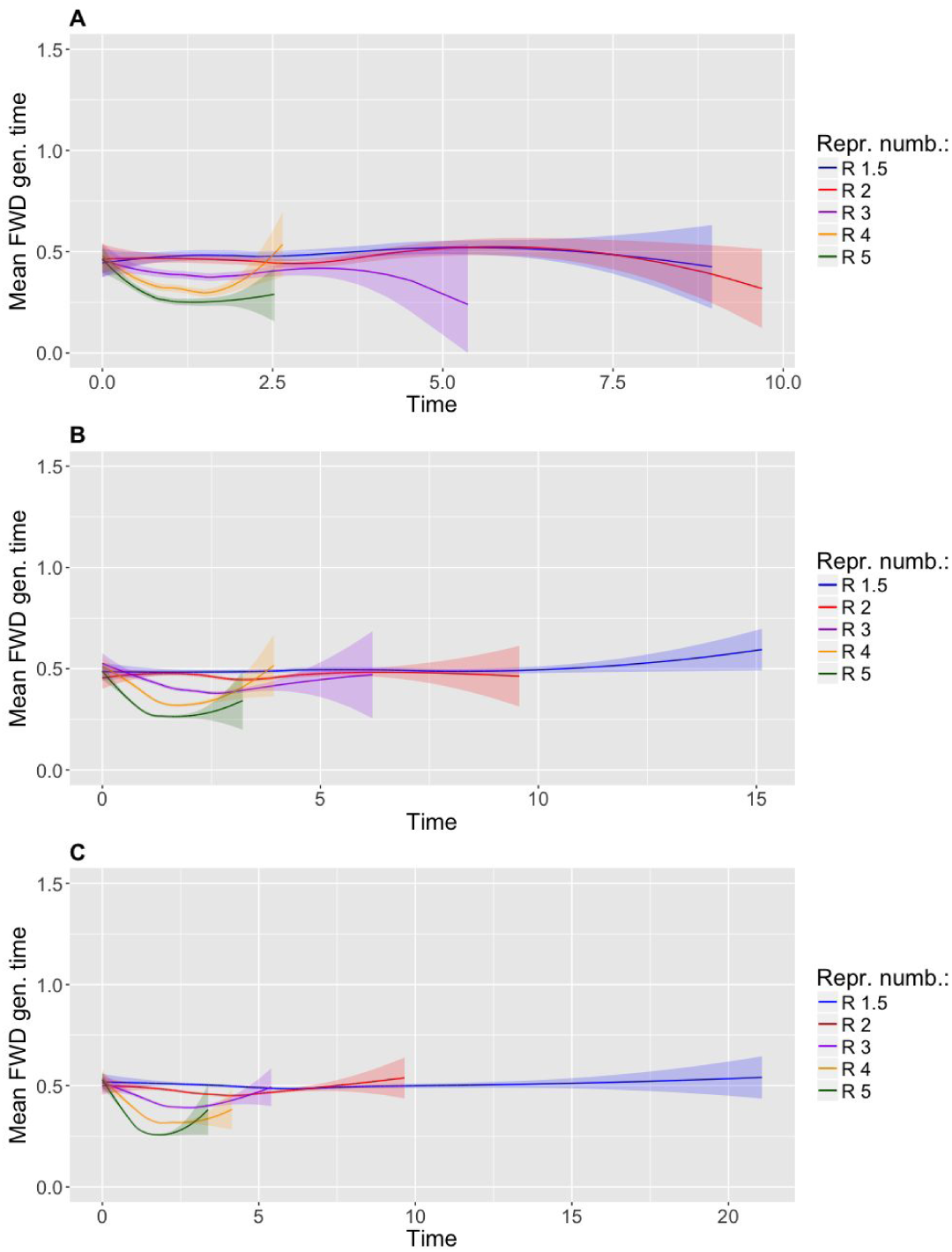
Evolution of the mean forward generation time - Population sizes. Evolution of the mean forward generation time in the baseline scenario for different population size. A, B and C identify, respectively, a population of size *N* = 100, 500, 1000. The infectious period considered is exponential with unitary rate and the reproduction number varies with *ℛ*_0_ = 1.5, 2, 3, 4, 5.

**S6 Fig.**
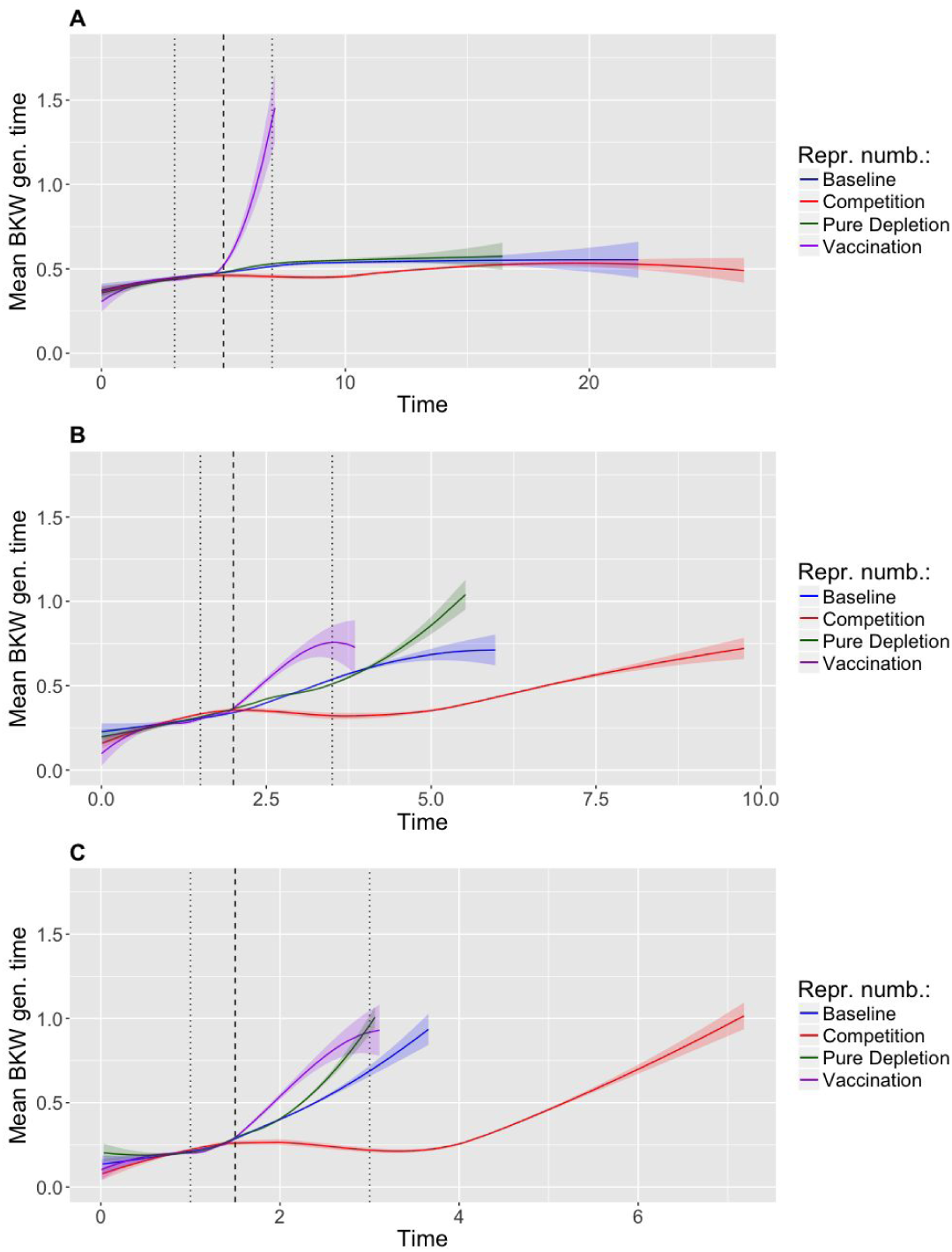
Evolution of the mean backward generation time. Evolution of the mean backward generation time comparing the baseline scenario, the vaccination (90% coverage) scenario, the competition scenario and the pure depletion scenario. The infectious period is constant, the population is of size *N* = 1000 and the considered reproduction number is, respectively, *ℛ*_0_ = 1.5 (A), *ℛ*_0_ = 3 (B) and *ℛ*_0_ = 5 (C).

